# A century of decline: loss of genetic diversity in a southern African lion-conservation stronghold

**DOI:** 10.1101/474940

**Authors:** Simon G. Dures, Chris Carbone, Andrew J. Loveridge, Glyn Maude, Neil Midlane, Ortwin Aschenborn, Dada Gottelli

## Abstract

**Aim:** There is a dearth of evidence that determines the genetic diversity of populations contained within present-day protected areas compared with their historic state prior to large-scale species declines, making inferences about a species’ conservation genetic status difficult to assess. The aim of this paper is to demonstrate the use of historic specimens to assess the change in genetic diversity over a defined spatial area.

**Location:** Like other species, African lion populations (*Panthera leo*) are undergoing dramatic contractions in range and declines in numbers, motivating the identification of a number of lion conservation strongholds across East and southern Africa. We focus on one such stronghold, the Kavango-Zambezi transfrontier conservation area (KAZA) of Botswana, Namibia, Zambia and Zimbabwe.

**Methods:** We compare genetic diversity between historical museum specimens, collected during the late 19^th^ and early 20^th^ century, with samples from the modern extant population. We use 16 microsatellite markers and sequence 337 base pairs of the hypervariable control region (HVR1) of the mitochondrial genome. We use bootstrap resampling to allow for comparisons between the historic and modern data.

**Results:** We show that the genetic diversity of the modern population was reduced by 12% to 17%, with a reduction in allelic diversity of approximately 15%, compared to historic populations, in addition to having lost a number of mitochondrial haplotypes. We also identify reduced allelic diversity and a number of ‘ghost alleles’ in the historical samples no longer present in the extant population.

**Main Conclusions:** We argue a rapid decline in allelic richness after 1895 suggests the erosion of genetic diversity coincides with the rise of a European colonial presence and the outbreak of rinderpest in the region. Our results support the need to improved connectivity between protected areas in order to prevent further loss of genetic diversity in the region.

## Introduction

Globally, mammal wildlife populations are reported to have undergone a 52% decline in the past half century (McRae et al., 2014), but over longer time periods the ranges and population declines have been far more severe (Ceballos, 2002; Janecka et al., 2014; Crees et al., 2016). While such studies focus on loses in population sizes and species’ distributions, relatively few have explored temporal losses in genetic diversity (Leonard, 2008), which may have significant impacts on a species’ ability to respond to environmental stochasticity and associated conservation interventions (Spielman et al., 2004).

Several reviews highlight insufficient genetic data available to decision makers as a major challenge in conservation genetics today (Frankham, 2010; Hoban et al., 2014). Genetic monitoring of individuals and populations over time was identified as one of the main topics in need of urgent attention. It is crucial to establish baseline genetic diversity measures against which future comparisons can be made to demonstrate decline or recovery (Jackson et al., 2011). To this effect the use of ancient museum samples provide an important genetic tool to measure within-species genetic diversity. This information in turn will be used to the development and implementation of strategies aim at minimizing genetic erosion and safeguarding genetic diversity.

One important flagship species that has undergone a major decline in population size and geographic range is the lion (*Panthera leo*) (Bauer et al., 2015a). Recent assessments of the lion population in Africa estimate between only 16,500 and 35,000 individuals remain (Riggio et al., 2013; Bauer et al., 2015b), with an estimated decline of 42% over the past 21 years (Bauer et al., 2015a; Bauer et al., 2015b). Major declines in wildlife populations across the region, however, have also been noted further back in time (Selous, 1908).

The dramatic decline of the African lion has made the protection of the remaining populations and the improvement of gene flow between populations of the upmost importance and has led to a number of transboundary conservation initiatives (Naidoo et al., 2012) such as the Kavango-Zambezi transfrontier conservation area (KAZA). The size of the KAZA region and its ability to support a large number of lion prides, results it being considered one of the few remaining strongholds for the lion population (Cushman et al., 2015). While this population, and the ability of lions to disperse long distances in the region, may be enough to sustain a robust population (Björklund, 2003), numbers do not necessarily allow us to understand all aspects of population status. Diminished populations are less effective at eliminating deleterious variants through selection (Xue et al., 2009; Spielman et al., 2004) making them vulnerable to reduced individual fitness, the loss of species’ evolutionary potential, and diminished ecosystem function and resilience. There is a risk of overestimating the potential for modern populations to resist the effects of demographic and genetic stochastic events on small populations if genetic factors are not considered. Populations which may be considered stable by contemporary conservation managers may in fact show signs of genetic erosion, thus needing greater conservation attention. However, currently there is no baseline genetic data for lion populations other than from the modern populations, which are likely to have also suffered major losses in genetic diversity (Björklund, 2003).

At the end of the 19^th^ and beginning of the 20^th^ century large numbers of animal specimens were being archived in natural history museums around the world (e.g. Dollman, 1921), including lions shot across the KAZA region. With the advent of methods to extract and analyse DNA from historic archival specimens (Higuchi et al, 1984; Leonard, 2008) there is the opportunity to assess the genetic diversity of populations pre-dating any significant anthropogenic influence. By comparing genetic data from museum collections with modern wild populations from the same area, one could assess the extent to which current levels of genetic variation have been reduced (Wandeler et al., 2007; Gebremedhin *et al.*, 2009).

To determine whether genetic diversity has declined over time, we compared genetic diversity between historical and modern lion populations from the KAZA region. We extracted DNA from historical lion samples taken from museum specimens in order to compare historic levels of genetic diversity against modern levels from the same region. We used a suite of microsatellite markers as well as sequencing of part of the hypervariable control region (HVR1) of the mitochondrial genome (mtDNA) to assess the degree to which genetic diversity in this population has been lost as a result of regional declines in lion numbers and distribution.

## Methods

### Samples

The Natural History Museum of London’s collections contain large numbers of lion skins and skulls from across the species range. The labelling of the collection data was of varying quality so specimens were cross-referenced with collector catalogues wherever possible. Twenty-seven lion specimens were sampled, originally collected from within the study region between 1879 until 1935 (Table 1, Fig. 1). Scrapes of any tissue remaining on the skulls or skin, or fragments of detached maxilloturbinal bone (thin bones inside the nasal cavities) were collected from each specimen.

**Table 1.**
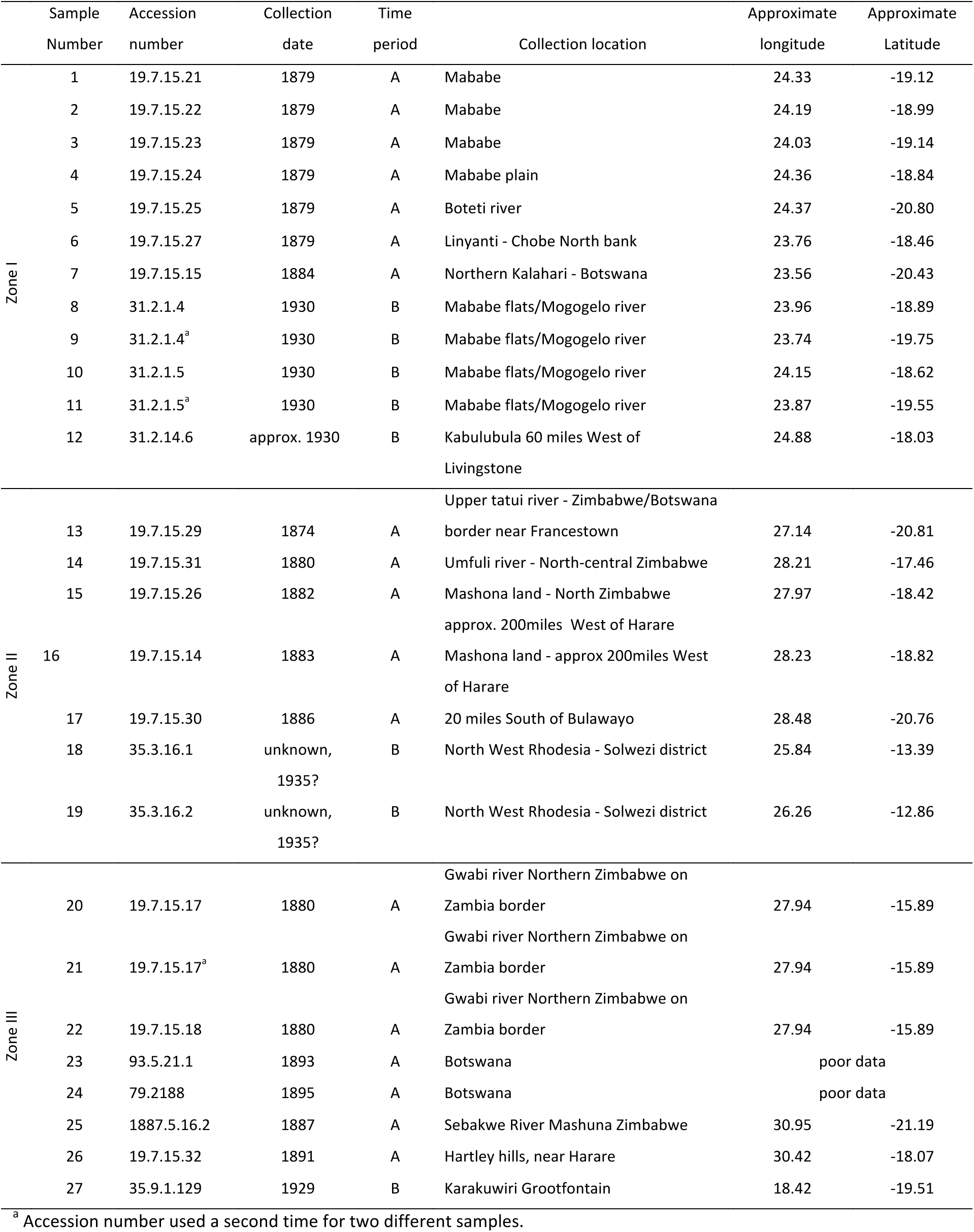
Museum samples from the Natural History Museum of London grouped according to three spatial zones. Sample Number refers to position on figure 1. Spatial zones represent; I) samples within the modern sampling area; II) the samples likely to be within maximum male dispersing distance of the modern sampling area, taken as 200km; III) all remaining samples across the region. Time periods represent samples collected between; A) 1874 to 1895; B) 1930-1935. Unclear dates use accession number as date reference. Longitude and latitude are estimated based on location data available for each specimen.

**Figure 1.**
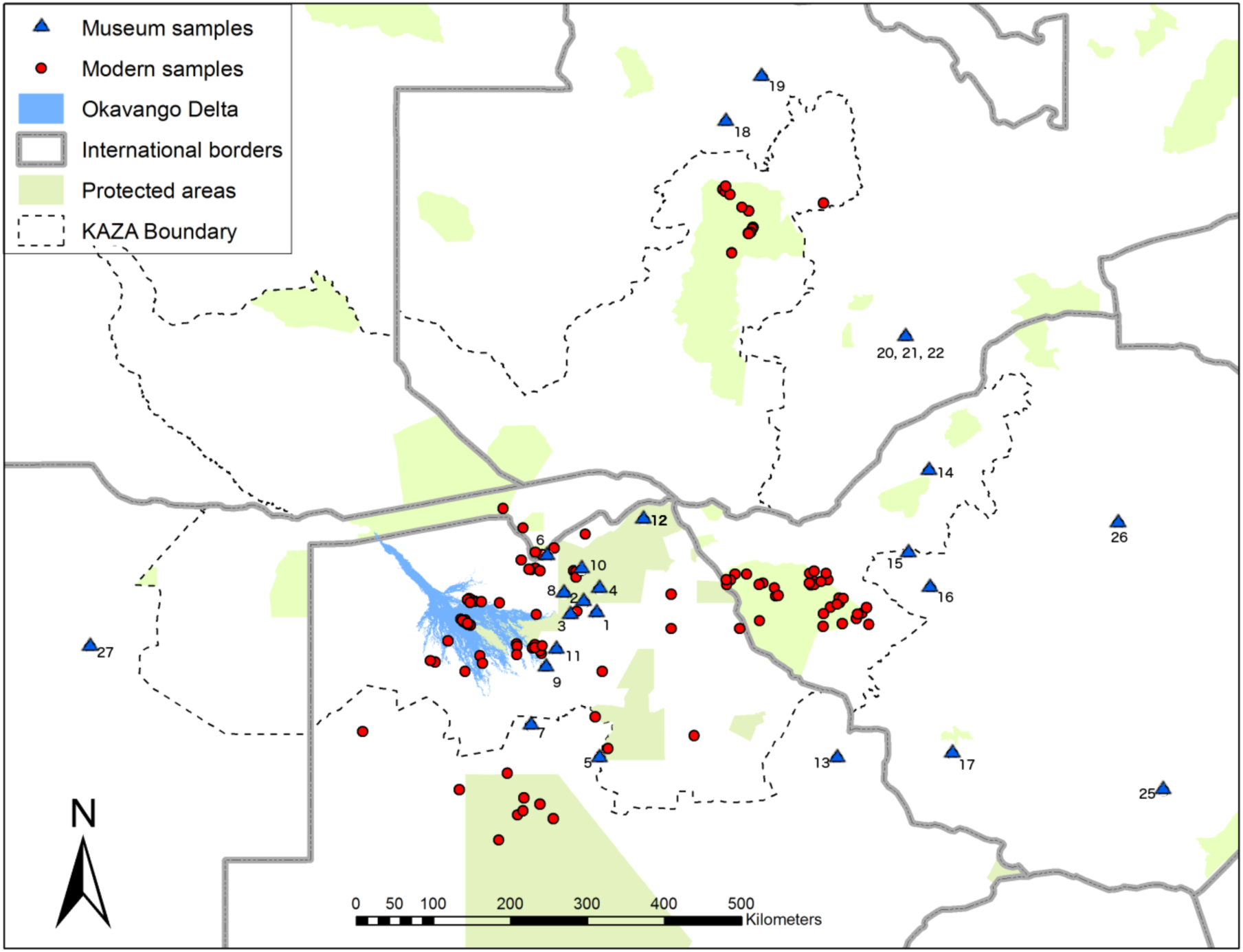
Map of Kavango-Zambezi region showing sampling distribution of modern lion samples (red circles) and museum samples (blue triangles and numbered)

Modern samples were collected from 204 free ranging wild lions between 2010 until 2013 (Fig. 1) in the form of blood (n=23), fresh tissue (n=113), dry tissue (n=13), faecal (n=14) and hair-pulls (n=41). Fresh tissue samples were collected using a remote biopsy dart system (Karesh et al., 1987). Hair pulls and blood were taken from immobilised animals. Dry tissue samples were taken from animals shot by the trophy hunting industry.

### Ancient DNA precautions

All pre-PCR work was performed in a laboratory exclusively devoted to ancient DNA, situated on a different floor from the PCR amplification laboratory and with an independent air handling system. To avoid sample cross-contamination a different set of equipment was used for each extraction (e.g. mortar and pestle, scalpel blades etc). Single-use equipment was immersed in sodium hypochlorite and removed from the working area after use. The working area was cleaned with sodium hypochlorite solution before work on the next sample commenced. All equipment was UV-irradiated overnight prior to further use. Filter tips were used to reduce cross contamination (Rohland & Hofreiter, 2007). Two blank extractions containing no tissue or bone were included during both extraction protocols to serve as negative extraction and PCR controls. Each fragment was independently amplified by PCR at least three times following the multi-tube approach (Taberlet et al., 1999) in an attempt to detect contamination and genotyping errors.

### DNA extraction

Total genomic DNA was extracted from each museum skin sample using approximately 25mg of tissue using DNeasy^®^ Blood and Tissue kits (Qiagen). We followed the manufacturer’s instructions but added a second incubation. To increase tissue lysis the first incubation was run overnight, for the second digestion we added a further 180µl Buffer ATL and 20µl proteinase K (600mAU/ml) and then incubated for a further 3 hours at 56°C.

DNA from bone samples was extracted using approximately 100mg of bone powder previously ground in a pestle and mortar. A master mix was prepared which, for each sample, comprised of 0.2ml 10% SDS (Invitrogen), 0.15ml proteinase K at 15mg/µl, a 1×1mm piece of DTT at 10mM and 1.65ml EDTA of pH 8.0 at 0.5M. This was warmed at 56°C until all ingredients dissolved and added to each bone-powder sample. Samples were incubated on a rotator at 56°C for 48 hours. Following digestion, tubes were centrifuged for one minute at 1300rpm and supernatant transferred to an Amicon^®^ Millipore Ultra Centrifuge filter which was centrifuged for 30 minutes at 1300rpm. A MinElute purification kit (Qiagen) was used to purify 100µl of extract following the manufacturer’s instructions, washing three times with PE buffer.

Modern DNA was extracted using approximately 25mg of tissue, 100μl of raw blood or 5-6 hair follicles using DNeasy^®^ Blood and Tissue kits (Qiagen) according to the manufacturer’s instructions. Faecal DNA was extracted using approximately 200mg of stool using QIAamp^®^ DNA Stool kits (Qiagen) according to the manufacturer’s instructions.

### Microsatellite amplification

We used sixteen microsatellite loci previously identified and amplified in the domestic cat (Menotti-Raymond et al., 1999) (FCA1, FCA45, FCA69, FCA75, FCA77, FCA96, FCA97, F115, FCA126, FCA129, FCA133, FCA193, FCA205, FCA224, FCA247, FCA391) which have previously been successfully used in lions (C. Driscoll, 1992; C. A. Driscoll et al., 2002; Dubach et al., 2013; Lyke et al., 2013; Spong & Creel, 2001). The nuclear marker primers were divided into multiplex combinations and fluoro-labelled with one of VIC, 6-FAM, PET or NED dyes, according to primer annealing temperatures and non-overlapping allele size range combinations (see Supplementary materials). See Supplementary Material for amplification conditions and sequencing details. The allele sizes and genotypes were scored in *GENEMAPPER v4.1* (Applied Biosystems).

### Mitochondrial sequencing

We amplified a 337bp hypervariable region (HVR1) of the *Panthera leo* mitochondrial control region, using previously published reverse and forward primers (Barnett et al., 2006). To improve the quality of the sequencing and avoid the problem of double banding due to the reverse primer being able to bind to multiple reverse sequence repeats, identified previously with these primers, we used a nested reverse primer designed for direct sequencing (Barnett et al., 2006). See Supplementary Information for PCR and sequencing conditions. Consensus sequence for each individual was obtained through alignment of the forward and reverse sequences in *GENEIOUS* (Kearse et al., 2012) to yield a minimum of 2x coverage for each base.

### Estimation of change in nuclear diversity

To detect changes in nuclear diversity between the modern and historic populations, using the microsatellite data, we calculated Nei’s unbiased estimate of expected heterozygosity (*H_E_*), observed heterozygosity (*H_O_*), inbreeding coefficient (*F_IS_*) and mean number of alleles per locus (*A*). This was performed using *GENEPOP* (Rousset, 2008) using methods documented in previous research on white-tailed eagles (Hailer et al., 2006). *GENEPOP* was also used to detect significant departures from Hardy-Weinburg equilibrium (HWE) and evidence of linkage disequilibrium within the sample groups. Unique alleles were identified for each time period using *CONVERT* (Glaubitz, 2004). The mean number of private alleles per locus found in each population was calculated using a rarefaction approach to control for differences in sample size, implemented in *ADZE* et al., 2008). DnaSp v.5 (Librado & Rozas, 2009) was used to calculate mtDNA haplotype diversity (*H*) and nucleotide diversity (π), as well as Tajima’s *D* to test for deviations from neutral evolution for both the modern and historic populations.

### Bootstrap resampling

There is an inherent inability to control the sampling design when using museum collections, including sample size, date and location of their collection. To allow comparisons between modern and historic nucleic diversity we used a bootstrapping procedure. When analysing the more rapidly mutating nuclear microsatellite data, we progressively restricted; *i*) the spatial extent of the historic samples, to match with more certainty the extent of the modern samples; *ii*) the time period over which the historic samples were collected, to restrict the possible influence of genetic drift with time within the sample set. Thus, we divided our historic data into three spatial zones representing; I) the samples within the modern sampling area; II) the samples likely to be within male dispersing distance of the modern sampling area, taken as 200km; III) all remaining samples across the region (Table 1). We also divided the historic data into two time periods, 1874-1895 (A) and 1929-1935 (B) (Table 1). The results from the historic samples sets were compared against our modern dataset using a bootstrapping procedure implemented in *POPTOOLS* (Hood, 2011). We created 100 populations of equal size to the historic data being used. Furthermore, to account for an apparent lack of historic sampling from within the Okavango Delta bootstrap sampling was repeated both with and without modern Okavango Delta samples. In a species such as lions, where female siblings tend to remain in the same pride or form a neighbouring pride and male siblings commonly forge a coalition, the likelihood of collecting data from close relatives was high. To test for the effects of close relatives, we followed the recommendations of Rodríguez-Ramilo & Wang (2012) and calculated all possible full-sibling and parent-offspring clusters in the programme *COLONY* (Wang & Scribner, 2014). We then randomly selected just one individual from each close-relative cluster, before re-rerunning the bootstrap procedure on the reduced data set.

### Mitochondrial ‘ghost’ alleles

Following the identification of all haplotypes present in the combined modern and historic data set, we were able to assess private haplotypes only present in one or other time period. Due to the much poorer quality of the museum sample data many sequences were considerably shorter than the modern counterparts, making direct comparisons of diversity difficult and lacking power. However, we were able to identify haplotypes only present in the historic data, likely to have been lost from the modern population (Leonard et al., 2005).

## Results

We achieved successful microsatellite amplification of all 27 museum samples and obtained usable mitochondrial sequences from 18 of these. A number of microsatellite loci could not be successfully genotyped across every sample, achieving a mean of 23.7 (s.d. ± 3.5) complete genotypes per locus (Data available on Figshare, DOI 10.6084/m9.figshare.3514469). No single locus or within group deviations from HWE were detected and tests for linkage disequilibrium were not significant after Bonferroni correction. Mitochondrial consensus sequence lengths varied from between 204 to 322bp, across a 337bp region (GenBank Accession no. KX661326 - KX661331).

In every bootstrap combination of our microsatellite data, regardless of how many samples were excluded, the historic lion population exhibited a higher heterozygosity, both observed (*t* = 8.75, *p* = 0.006) and expected (*t* = 14.80, *p* = 0.002). The same results for reduced heterozygosity were returned when the Okavango lions were removed from the analysis (observed, *t* = 8.75, *p* = 0.006; expected, *t* = 14.79, *p* = 0.002).

In every iteration of the data the modern population showed a much greater deficiency in the observed heterozygosity compared to the expected, represented by a significantly larger inbreeding coefficient (*F_IS_*) for all modern sample sets (*t* = 5.42, *p* = 0.016; Table 2). The reduction in the geographic extent of the historic data resulted in a limited change in the observed heterozygosity from 0.7565 for the broadest sample set, up to 0.7975 for the most limited. When we control for differences in sample size (n=27 vs. 12) using 100 bootstrap replications the observed heterozygosity for the full sample set of zones I-III increased from 0.7565 to 0.7612, similar to levels observed among the more spatially restricted data encompassing just zones I and II.

**Table 2.** Genetic diversity for the Kavango-Zambzi African lion population within each spatial scale for 16 microsatellite loci. Modern samples represent the average value from 100 bootstrap replications including or excluding the Okavango samples respectively. *N* = sample size; *H_E_* = expected heterozygosity *H_O_* = observed heterozygosity; *F_IS_* = inbreeding coefficient; *A* = mean number of alleles per locus; s.d. = standard deviation of bootstrap replications.

| | Sample set | $N$ | $H_E$ | s.d. | $H_O$ | s.d. | $F_{IS}$ | $A$ | s.d. |
| --- | --- | --- | --- | --- | --- | --- | --- | --- | --- |
| Zone I-III | Historic | 27 | 0.7813 | - | 0.7565 | - | 0.032 | 8.50 | - |
|  | Modern | 27 | 0.6989 | (0.014) | 0.6541 | (0.025) | 0.064 | 6.55 | (0.37) |
|  | Modern - without Okavango | 27 | 0.7186 | (0.013) | 0.6688 | (0.020) | 0.069 | 7.00 | (0.37) |
| Zone I-II | Historic | 19 | 0.7807 | - | 0.7676 | - | 0.017 | 7.69 | - |
|  | Modern | 19 | 0.6928 | (0.017) | 0.6483 | (0.025) | 0.064 | 6.23 | (0.35) |
|  | Modern - without Okavango | 19 | 0.7169 | (0.014) | 0.6647 | (0.021) | 0.073 | 6.49 | (0.35) |
| Zone I | Historic | 12 | 0.7945 | - | 0.7975 | - | -0.004 | 6.75 | - |
|  | Modern | 12 | 0.6946 | (0.023) | 0.6523 | (0.034) | 0.061 | 5.39 | (0.30) |
|  | Modern - without Okavango | 12 | 0.7146 | (0.021) | 0.6606 | (0.035) | 0.076 | 5.60 | (0.34) |

Across the data we identified 29 alleles present only in the historic samples and 54 private alleles only found in the modern data, however the latter come from a much larger data set. The mean number of private alleles is consistently higher in the historic data than in the modern data when controlling for sample size (Fig. 2). Such ‘ghost alleles’ (Bouzart et al., 1998; Groombridge et al., 2000) were identified in 14 out of the 16 microsatellite markers, only absent from Fca126 and Fca391. Even when reducing the historic data to only those within the most conservative spatial area (n=13) we still found 18 alleles not present in the modern samples, spread across all microsatellite markers except Fca126, Fca129, Fca193 and Fca391.

**Figure 2.**
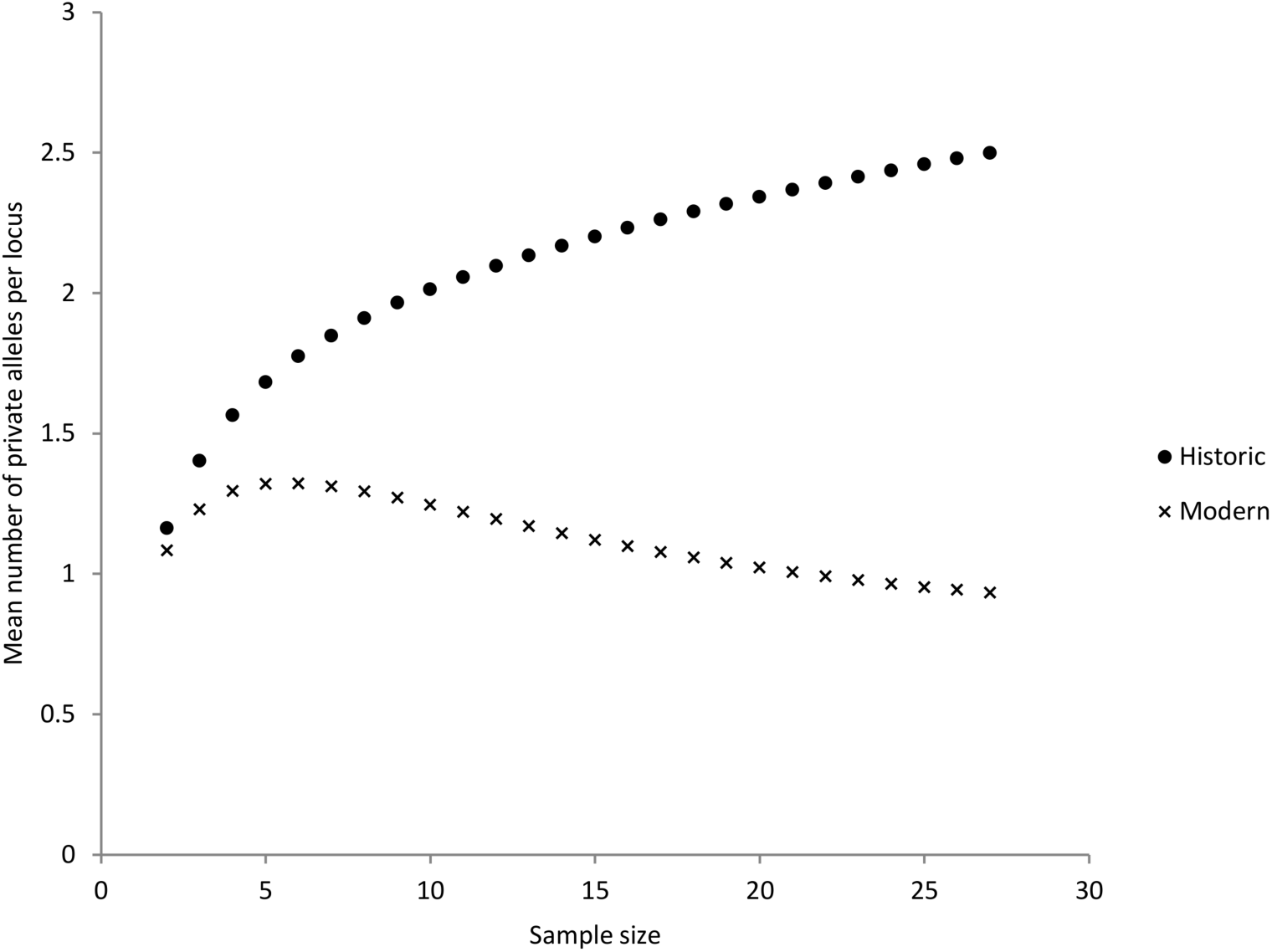
Mean number of private alleles per locus as a function of standardised sample size for historic and modern microsatellite data.

Removing samples collected between 1929-1935 made no difference to heterozygosity across the data (see Supplementary materials), however, it did result in a reduction in the allelic richness from 7.5 to 6.29, the latter being similar to the present day values. When we reduced the data to include only samples collected between 1929-1935, the allelic richness (5.88) closely matches that found within the modern samples.

Removing close relatives had a negligible effect on any values. In the full modern data set the observed heterozygosity increased from 0.6541 to 0.6570, expected heterozygosity from 0.6989 to 0.7039, the inbreeding coefficient from 0.064 to 0.066 and the mean number of alleles from 6.55 to 6.65.

The mtDNA data (Table 3) indicates six haplotypes present within the historic dataset (*H* = 0.6993, *π* = 0.00065), but three of these appear to be missing from the extant lions (*H* = 0.3257, *π* = 0.0007). Tajima’s *D* for both the historic (*D* = −1.09629; *p* < 0.1) and modern (*D* = −0.53568, *p* < 0.1) population are negative but not significant, suggesting no deviation from neutrality. Aside from the three ‘ghost’ haplotypes identified, there may be others present within the same mtDNA region that due to the degradation of the historic DNA remain unidentified. Since two of the ‘ghost’ haplotypes were identified from single individuals, each only with a single nucleotide insertion, we must caution that they may be false haplotypes caused by DNA degradation (Wandeler et al., 2007). Even following a more conservative approach, one previously common haplotype remains unrepresented in the modern samples.

**Table 3.** Mitochondrial DNA control region haplotypes from historical specimens and the extant lion population of the KAZA region. “-“ and “N/A” represent a deletion or missing sequence data, respectively, at the specified nucleotide position.

| Sample size |  | Haplotype | Variable nucleotide position<sup>a</sup> |  |  |  |  |
| --- | --- | --- | --- | --- | --- | --- | --- |
| Modern | Historic |  | 221 | 343 | 367 | 368 | 378 |
| 31 | 5 | *i* | - | - | T | A | - |
|  | 9 | *ii* | T | - | T | A | - |
|  | 1 | *iii* | N/A | - | T | A | C |
|  | 1 | *iv* | N/A | C | T | A | - |
| 3 | 1 | *v* | - | - | C | A | - |
| 4 | 1 | *vi* | - | T | T | G | - |

<sup>a</sup> 1 corresponds to position 16,176 in the complete *Panthera leo leo* mtDNA sequence (Ma & Wang, 2014).

## Discussion

The value of genetic diversity is increasingly recognized for contributing to individual fitness, species’ evolutionary potential, and ecosystem function and resilience (Whitham et al. 2008). There is therefore an urgent need for policy-relevant studies to help define sensitive and robust indicators of changes in genetic diversity (Hoban et al. 2013a).

Our analysis demonstrates that over the past century the lion population of the Kavango-Zambezi region has lost genetic diversity. Contemporary observed heterozygosity has been reduced by 12% to 17% compared to historic populations. Despite having a number of missing alleles across the samples, genetic diversity was still historically higher than in the contemporary lion population. The decline in heterozygosity is not as dramatic as that seen in some highly threatened or bottlenecked species, for example, 57% in the Mauritius kestrel (*Falco punctatus*) (Groombridge et al., 2000) or 43% in sea otters (*Enhydra lutris*) (Larson et al., 2002), it nevertheless represents a worrying reduction in diversity considering this population is one of only six lion strongholds remaining in Africa.

While the low sample size of the bootstrapping means caution should be taken before extrapolating to the true *F_IS_*, it is clear that the reduced heterozygosity exposes lions of the region to a higher risk of inbreeding depression than their historic counterparts. As well as clear decline in nuclear diversity, as assessed with the microsatellite analysis, there is also an indication of a loss in mitochondrial diversity. One haplotype detected in multiple historic samples, and two more haplotypes detected in single samples, remain entirely undetected in the modern population. The results are in agreement with previous research which has identified both declining populations and increasing fragmentation in the region (Elliot et al., 2014; Loveridge et al., 2007).

Similar to other species, the global decline in lion numbers has largely been driven by human-wildlife conflict and habitat loss (Keyghobadi et al., 2005; Bauer et al., 2015b). Given the rapid expansion of human activities in the region in the 20^th^ century, the downward trend in genetic diversity we observed is perhaps unsurprising and seemingly confirms the pessimistic observations made in the late 19^th^ century. For example, one account from Frederick Courtney Selous records, “During the twenty years since my first arrival in 1871, I…had seen game of all kinds gradually decrease and dwindle in numbers to such an extent that I thought that nowhere south of the Great Lakes could there be a corner of Africa left where the wild animals had not been very much thinned out” (Selous, 1908). Interestingly, allelic richness did not differ between the intermediate temporal (1929-1935) and contemporary population samples, suggesting that allelic richness was lost prior to the intermediate sampling period. A temporal decline in genetic diversity of the historic samples was not detected through measures of heterozygosity, likely due to changes in allelic richness being detectable before population declines impact upon heterozygosity (Athrey et al., 2011). The rapid decline observed in allelic richness does coincide with the arrival of the first western settlers in 1890 and the subsequent rise of the colonial presence in the region after the end of the Matabele Wars in 1897 (Parsons, 1993). Furthermore, modern firearms became more prevalent following European settlement and predators were often persecuted as vermin (Woodroffe, 2000), which likely contributed to the earlier decline of lions in the study region. Whilst the timing of genetic decline and colonial settlement is compelling enough to suggest causation, the evidence is not conclusive.

The epizootic of the rinderpest virus also struck during the late 1890’s resulting in the death of vast populations of buffalo, giraffe and wildebeest, as well as domestic livestock (Van den Bossche et al. 2010). Such an epidemic is very likely to have also had a considerable impact on the predators of the region.

Given the level of habitat loss and fragmentation observed across sub-Saharan Africa (Keyghobadi et al., 2005; Bauer et al., 2015b), the increased threat of epizootics facilitated by human movements (Butler et al., 2004), as well as the impacts of a changing climate (Thomas et al. 2004), it is imperative that efforts are made to conserve genetic diversity. Without such genetic diversity, a species resilience and ability to adapt to future stochastic events becomes greatly compromised (Whitham et al. 2008). This study provides quantitative data on temporal genetic monitoring that is urgently needed to optimize conservation and management efforts. Since KAZA is considered one of the more stable lion populations in Africa, the work presented here should provide motivation for increased conservation action to safeguard against further loss of genetic diversity of lions and other species across the region (Krofel et al., 2015). In particular greater connectivity between lion population in protected areas across the region and thus the mixing of genetic material should be supported (Cushman et al., 2015).

## Data Accessibility Statement

Microsatellite data is available at Figshare, DOI 10.6084/m9.figshare.3514469

Mitochondrial sequence data has been submitted to the GenBank database under accession no. KX661326 - KX661331.

## Acknowledgements

We thank NHM London for access to historic samples; the Botswana Department of Wildlife and National Parks for providing research and collection permits (EWT 8/36/4 XIII [35]); Debbie Peak, Rob Jackson, Kyle burger, Robyn Coetzee, Robert Riggs & Botswana Predator Conservation Trust for contributing samples; the staff at Wilderness Safaris Botswana, Chitabe, and Machaba; Crispin Sanderson, Grant Huskisson, Dane Hawk, Rick Nelson, Erik Verreynne, Alan Wilson, Anna Butterfield & Jaques Van deMerwe of Vision International, Wilton Raats, Dominik Bauer and Kristina Kesch for logistical and veterinary support; Anton van Schalkwyk and Hanri Ehlers for invaluable support and funding; PneuDart for equipment; Wilderness Wildlife Trust for financial support. We also thank Jinliang Wang for helpful comments on previous drafts. All import and export permits were granted. SGD was supported by a BBSRC CASE-studentship (BB/F017324/1).

